# Exploring the mechanism of Panax Notoginseng in the treatment of skin wound based on network pharmacology and experimental verification

**DOI:** 10.64898/2026.02.28.708691

**Authors:** Yi-bin Li, Qing-lin Li, Jun Liu, Jun-cen Li, Hua-man Geng, Gen-ke Li, Chen Jin, Jie Luo, Zhi-hong Zhang

## Abstract

**Background:** When the skin wound defect is too large, it is difficult for the body to heal itself, and medical treatment is often needed. How to shorten the healing cycle and reduce the incidence of infection is a difficult problem faced by clinicians. Panax notoginseng(PN), a traditional Chinese medicine, can promote the absorption of inflammatory exudates, granulation tissue formation and epidermal proliferation, effectively inhibit the inflammatory reaction of wounds and promote the healing of skin wounds, but its molecular mechanism has not been fully clarified so far. Based on network pharmacology and animal experiments, this study explored the target and molecular mechanism of PN in the treatment of skin wound.

**Methods:** The active components and potential targets of PN were obtained from the Traditional Chinese Medicine System Pharmacology Database and Analysis Platform (TCMSP) and UniProt database, and the skin wound-related targets were obtained from the GeneCards database. The intersecting targets were filtered using Venny 2.1.0. The intersecting targets were imported into the STRING database to construct a protein-protein interaction (PPI) network. Cluster analysis was performed using the MCODE and CytoHubba plugins in Cytoscape 3.8.2 to obtain core functional network modules and the top 10 key target genes. The intersecting targets were subjected to KEGG and GO enrichment analysis using the DAVID 6.8 database (https://david.ncifcrf.gov/). The component-target-pathway network of PN in the treatment of skin wounds was constructed using Cytoscape 3.8.2 software. In the experimental verification phase, 48 Sprague-Dawley (SD) rats were randomly divided into a control group and a PN group, with 24 rats in each group. A full-thickness skin excision was performed to establish a wound model. The PN group received intraperitoneal injections of the drug, while the control group received an equivalent amount of saline. Wound area measurements were taken on days 1, 4, and 7 after model establishment. Histopathological changes in the injured area and the expression and localization of TNF-α, IL-6, and IL-10 were observed through hematoxylin and eosin (HE) staining and immunohistochemical staining. Relative expression levels of the three factors were detected using quantitative real-time polymerase chain reaction (qRT-PCR) and enzyme-linked immunosorbent assay (ELISA).

**Results:** This study identified 8 major active components, 156 targets, and 115 signaling pathways involved in the treatment of skin wounds in rats using PN. The top 10 core target genes included TNF, IL-6, and IL-10, primarily enriched in signaling pathways such as NF-κB, MAPK, and JAK-STAT. Animal experiments revealed that at 4 and 7 days post-injury, the wound area in the PN group was significantly smaller than that in the control group (P<0.05). HE staining showed reduced infiltration of neutrophils and inflammatory cells in the injury area at 7 days in the PN group, accompanied by more pronounced fibroblast proliferation and collagen secretion. Molecular detection indicated that TNF-α, IL-6, and IL-10 positive reactants were mainly distributed in the cytoplasm and matrix of epidermal cells, inflammatory cells, and fibroblasts in the skin. qRT-PCR and ELISA results showed that TNF-α expression in the PN group was significantly lower than that in the control group at 4 and 7 days (P<0.01). IL-6 expression was lower than that in the control group at all time points, peaking at 4 days and then decreasing (P<0.01). IL-10 expression was significantly lower than that in the control group at 1 and 7 days (P<0.01).

**Conclusion:** PN exhibits characteristics such as multi-component, multi-target, multi-pathway synergy, and multiple regulatory pathways in the treatment of skin wounds. It can reshape the dynamic balance of cytokine networks, including TNF-α, IL-6, and IL-10, optimize the temporal progression of “inflammation initiation - repair transition - tissue remodeling”, and provide a therapeutic effect of “efficient debridement - orderly repair - low scar risk” for skin wounds. It is one of the ideal natural drugs for regulating skin wound healing.

## Background

Skin wound repair has long been a subject of significant interest. When skin defects are too large, the body struggles to heal them spontaneously, often requiring medical intervention for treatment [1]. The wound healing process comprises the hemostasis phase, inflammatory response phase, proliferative phase, and tissue remodeling phase [2]. During the inflammatory response phase, the body releases large quantities of pro-inflammatory cytokines such as tumor necrosis factor-α (TNF-α), interleukin-6 (IL-6), interleukin-1 (IL-1), and gamma interferon (IFN-γ). Concurrently, anti-inflammatory cytokines like interleukin-10 (IL-10), transforming growth factor-β (TGF-β), IL-4, and others. A dynamic equilibrium exists between these two groups. Inflammatory factors activate local and systemic cascade inflammatory responses to resist pathogen invasion. However, when skin defects are too large, the imbalance in inflammatory factor secretion often leads to uncontrolled inflammatory reactions. This prolongs wound healing time and makes the body more susceptible to pathogenic bacteria, triggering wound infection [3]. Shortening the healing cycle and reducing infection incidence remain significant challenges for clinicians.

Panax notoginseng(PN) belongs to the Araliaceae family. It tastes sweet, slightly astringent, and slightly bitter, with a warm nature, and is attributed to the liver, stomach, and large intestine meridians [4]. Its chemical components include saponins, flavonoids, volatile oils, amino acids, polysaccharides, starch, proteins, notoginsenoside, and various trace elements [5]. Currently, Panax notoginseng has been formulated into various dosage forms such as tablets and injections for clinical use. Among these, the commonly used trade name for Panax notoginseng injection is Xuesaitong injection. According to literature reports [6, 7], PN can promote the absorption of inflammatory exudate from wound surfaces, facilitate the formation of granulation tissue, and stimulate epidermal proliferation. It effectively inhibits wound inflammation and promotes skin wound healing. Although there are numerous studies on the treatment of skin wounds using single or compound Chinese herbal medicines, most of them are empirical summaries and still lack exploration of the therapeutic mechanisms. Therefore, this study employs network pharmacology analysis and establishes a rat skin wound repair model for experimental verification, aiming to further explore the potential molecular mechanisms of PN in treating skin wounds. This provides new insights for clinical treatment of skin wounds and also offers a reliable theoretical basis for the drug development of PN.

## Methods

### Core target screening and signal pathway analysis

TCMSP database (http://tcmspw.com/) [8] was used to set the pharmacokinetic screening conditions: oral bioavailability (OB) ≥ 30% and drug-like property (DL) ≥ 0.18 [9, 10]. Obtain that active compound of the traditional Chinese medicine Notoginseng and the corresponding target. The GeneCards database (https://www.genecards.org/) was used to collect the target of skin trauma, and the target with Score value > median was selected as the potential target of skin trauma in the database. Then the above targets are proofread and duplicated in UniProt database (http://www.uniprot.org/uploadlists/). Venny2.1.0 website (https://bioinfogp.cnb.csic.es/tools/venny/) was used to intersect the target points of notoginseng and skin trauma, and the intersection targets were submitted to STRING database (https://string-db.org/) to construct PPI network. Using MCODE and CytoHubba plug-ins in CytoScape 3.8.2 for cluster analysis, the core functional network module of interaction between targets and the top 10 key target genes were obtained. The enrichment analysis of KEGG and GO was carried out by using DAVID 6.8 database (https://david.ncifcrf.gov/). The component-target-pathway network of PN was constructed by Cytoscape3.8.2 software.

### Animals and treatment

Forty-eight 2-month-old SD male rats, weighing 180-200 g with an average of (180±20) g, were provided by Shanghai Jiagan Biotechnology Co., Ltd., The animal experimental plan was approved by the Experimental Animal Ethics Committee of HuangHuai University (Ethics approval number: 202403170027). SD rats were randomly divided into PN group (experimental group) and control group. Before the experiment, the rats were shaved, weighed, anesthetized by intramuscular injection with Zoletil (Batch No.: 83887905) (dose: 0.5 ml/kg), fixed on the operating table, and routinely disinfected; A circular punch with a diameter of 1cm was used to punch holes at the skin 0.5 cm away from the spine without damaging the fat and fascia on the muscles adjacent to the spine. After successful modeling, the rats were reared in a single cage. The medication was started 2 hours after operation. The PN group was injected intraperitoneally with Panax notoginseng (dose: 1.5 ml/animal), and the control group was injected intraperitoneally with the same dose of normal saline. Once a day for 1 week, standard feed was given, clean water was supplied, the feeding temperature was maintained at 25 °C, and the bedding of the squirrel cage was changed every 1 day. Materials were collected at 1d, 4d and 7d after operation. Before the samples were collected, Zoletil was injected intramuscularly for anesthesia, and the whole-thickness skin tissue within 5mm of the wound edge was cut. Some of them were fixed in 4% formaldehyde solution, and some were stored in -80°C refrigerator for later use.

### Main reagents

Panax notoginseng injection(Xuesaitong Injection) was purchased from Yunnan phytopharmaceutical Co., Ltd., with national medical products administration approval number Z53020135 and the specification of 2ml: 100mg. Interleukin-6(IL-6) monoclonal antibody (Article No.: 66146-1-Ig, Proteintech Group,lnc.), Tumor Necrosis Factor -α(TNF-α) monoclonal antibody (Article No.: 60291-1-Ig, Proteintech Group,lnc.), Interleukin-10 (IL-10) monoclonal antibody (Article No.: 60269-1-Ig, Proteintech Group, Inc.). Enzyme linked immunosorbent assay (ELISA) kit for Rat IL-6 (Article No.: DY506), Rat IL-10 ELISA kit (Article No.: KE200003) and Rat TNF-α ELISA kit (Article No.: KE20001).

### Hematoxylin eosin (HE) staining

1). Sample fixation: Tissue sections were baked at 60 °C for 3-4 hours; 2). Dewaxing: 3 xylene treatments for 10-15 minutes each; 3). Hydration: 100%, 95%, 85% and 75% gradient alcohol treatment sequentially, each stage is 2 minutes, followed by running water rinsing for 5 minutes; 4). Hematoxylin staining: stain for 15-20 minutes, rinse with running water for 1 minute; It was differentiated by 1% hydrochloric acid alcohol for 3 seconds, and then the running water returned to blue for 45 minutes; 5). Eosin staining: staining with acidified eosin ethanol solution for 2 minutes; 6). Dehydration transparency: dehydrated by 75%, 85%, 95% and 100% gradient alcohol, and then treated with xylene for 3 times, each time for 10 minutes; 7). Sealing observation: After sealing with neutral resin, the dyeing quality and sample structure were evaluated under a microscope.

### Immunohistochemical(IHC) staining

1). Dehydration: Paraffin sections are treated with xylene I and II for 15 minutes each → anhydrous ethanol I and II for 5 minutes each → 85% and 75% ethanol for 5 minutes each → rinsed with distilled water; 2). Antigen retrieval: Heat citric acid buffer (pH6.0) in a microwave oven (to prevent evaporation of the solution and drying of the slides). After cooling, decolorize with PBS and wash on a shaker for 3 times (5 minutes each time); 3). Blocking endogenous peroxidase: Incubate with 3% hydrogen peroxide at room temperature for 25 minutes (in the dark), then wash three times with PBS (pH 7.4); 4). Blocking and primary antibody incubation: After blocking with BSA/serum, add diluted primary antibody dropwise and incubate overnight in a humidified box at 4°C; 5). Secondary antibody incubation and color development: After washing with PBS, add HRP-labeled secondary antibody and incubate at room temperature for 50 minutes. Develop with DAB (positive color is brownish yellow), and rinse with tap water to stop the reaction; 6). Counterstaining and bluing: Harris hematoxylin counterstaining → hydrochloric acid alcohol differentiation → ammonia water bluing → rinsing with running water; 7). Dehydration and mounting observation: After gradient dehydration, mount the slides with neutral paraffin, observe them under a microscope, and perform image scanning and analysis.

### Real-time quantitative PCR detection(qRT-PCR)

The relative mRNA expression levels of TNF-α, IL-6, and IL-10 in each group of rats were detected using qRT-PCR. Total RNA was extracted using TRIzol reagent (Invitrogen). The extracted mRNA was reverse transcribed into cDNA using Promega’s MMLV system. The cDNA served as the template for real-time PCR experiments. The reaction system was prepared according to the instructions provided in the dye-based PCR master mix kit (TaKaRa). The reaction conditions were as follows: 95°C for 1 minute (preheat) -- 95°C for 15 seconds -- 60°C for 30 seconds (40 cycles) - - melting curve (95°C for 15 seconds -- 60°C for 1 minute -- 95°C for 15 seconds). The Q-PCR results were calculated using the 2-ΔΔCt method to determine the relative expression levels of each target mRNA. The primer information for each gene was designed by Daixuan Biotechnology and synthesized by Invitrogen. See **Table 1** for details.

**Table 1.**
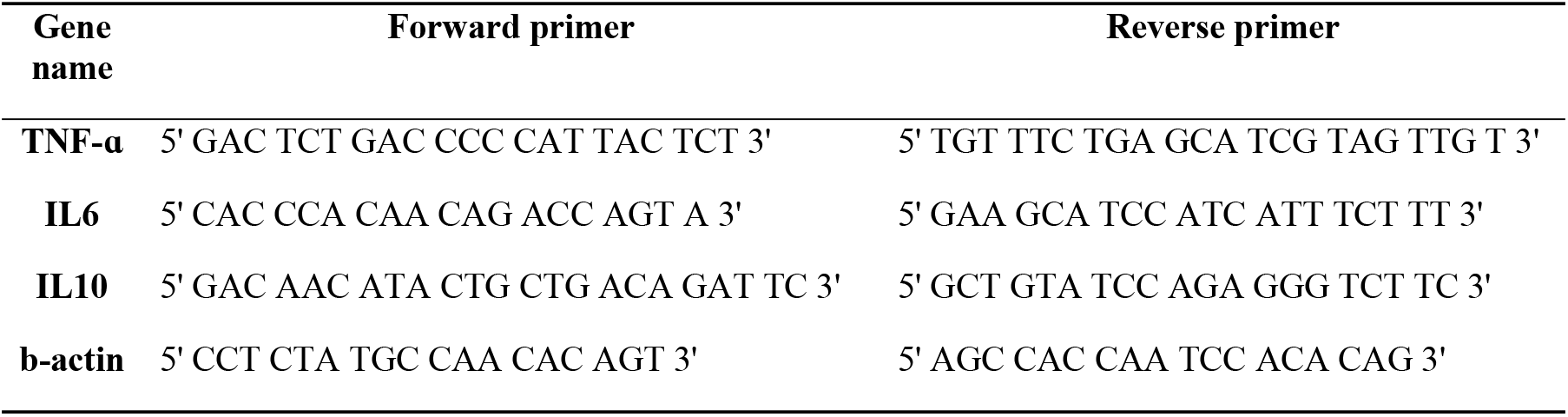
Primer sequence information.

### ELISA detection

Tissue samples from the skin injury areas of rats in both the PN group and the control group were collected. The tissues were washed with pre-cooled PBS (0.01M, pH 7.4) to remove any residual blood or impurities on the surface. The tissue pieces were weighed and cut into as small fragments as possible. An appropriate amount of pre-cooled PBS (at a weight-to-volume ratio of 1:9, with 1g of tissue sample corresponding to 9mL of PBS) was added, and the samples were homogenized thoroughly using an electric homogenizer on ice. The homogenate was then subjected to ultrasonic disruption to further lyse the tissue cells. The homogenate was aspirated into centrifuge tubes and centrifuged at 5000×g for 5 minutes at 2-8°C. The supernatant was collected and stored at -20°C for future use, avoiding repeated freeze-thaw cycles. According to the instructions provided with the ELISA kits for TNF-α, IL-6, and IL-10, the levels of each indicator in the rat skin trauma area were detected.

### Statistical analysis

After successful modeling, images were captured using a Sony digital camera, and the wound area of the rat skin was measured using Image-Pro Plus 6.0 software. Tissue section images were captured using an OLYMPUS digital camera. The data were expressed as mean ± standard deviation 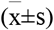. SPSS 20.0 software was used for inter-group comparisons using independent sample t-tests, and for intra-group comparisons using the LSD method of analysis of variance. A p-value < 0.05 was considered statistically significant.

## Results

### Target protein screening and interaction

This study obtained 8 potential active components of PN from the TCMSP database (**Table 2**), along with their corresponding 156 possible targets. From the GeneCards database, using “skin wound” as the keyword, 5,128 skin wound targets were identified. Targets with a Score value greater than the median were set as potential targets for skin wound. The maximum Score value was 60.05, the minimum was 0.15, and the median was 2.06. Therefore, targets with a Score value of 2.06 or above were considered potential targets for skin wound, ultimately resulting in 2,565 potential targets. A Venn diagram was created using venny2.1.0, revealing 124 intersecting targets between PN and skin wound (**Fig 1A**). These targets were submitted to the STRING platform, with the species set to “Homo sapiens” and the clustering method set to “kmeans clustering”, to obtain the PPI network interaction diagram of the intersecting targets (**Fig 1B**). The MCODE plugin in CytoScape 3.8.2 software was used to perform cluster analysis on the interaction relationships, resulting in the highest-scoring core subnetwork (**Fig 1C**). The cytoHubba plugin was applied to obtain the top 10 key genes ranked by Maximal Clique Centrality (MCC) analysis (**Fig 1D**), which were: tumor necrosis factor (TNF), interleukin-6 (IL-6), C-C motif chemokine ligand 2 (CCL2), interleukin-1β (IL1B), interleukin-1α (IL1A), C-X-C motif chemokine ligand 8 (CXCL8), interleukin-10 (IL10), C-X-C motif chemokine ligand 10 (CXCL10), Jun proto-oncogene (JUN, AP-1 transcription factor subunit), and RELA proto-oncogene (RELA, NF-KB subunit).

**Table 2.** Main active components of Panax notoginseng.

| Mol ID | Compound | OB% | DL |
| --- | --- | --- | --- |
| MOL001494 | Mandenol | 42 | 0.19 |
| MOL001792 | DFV | 32.76 | 0.18 |
| MOL002879 | Diop | 43.59 | 0.39 |
| MOL000358 | beta-sitosterol | 36.91 | 0.75 |
| MOL000449 | Stigmasterol | 43.83 | 0.76 |
| MOL005344 | ginsenoside rh2 | 36.32 | 0.56 |
| MOL007475 | ginsenoside f2 | 36.43 | 0.25 |
| MOL000098 | quercetin | 46.43 | 0.28 |

**Fig 1.**
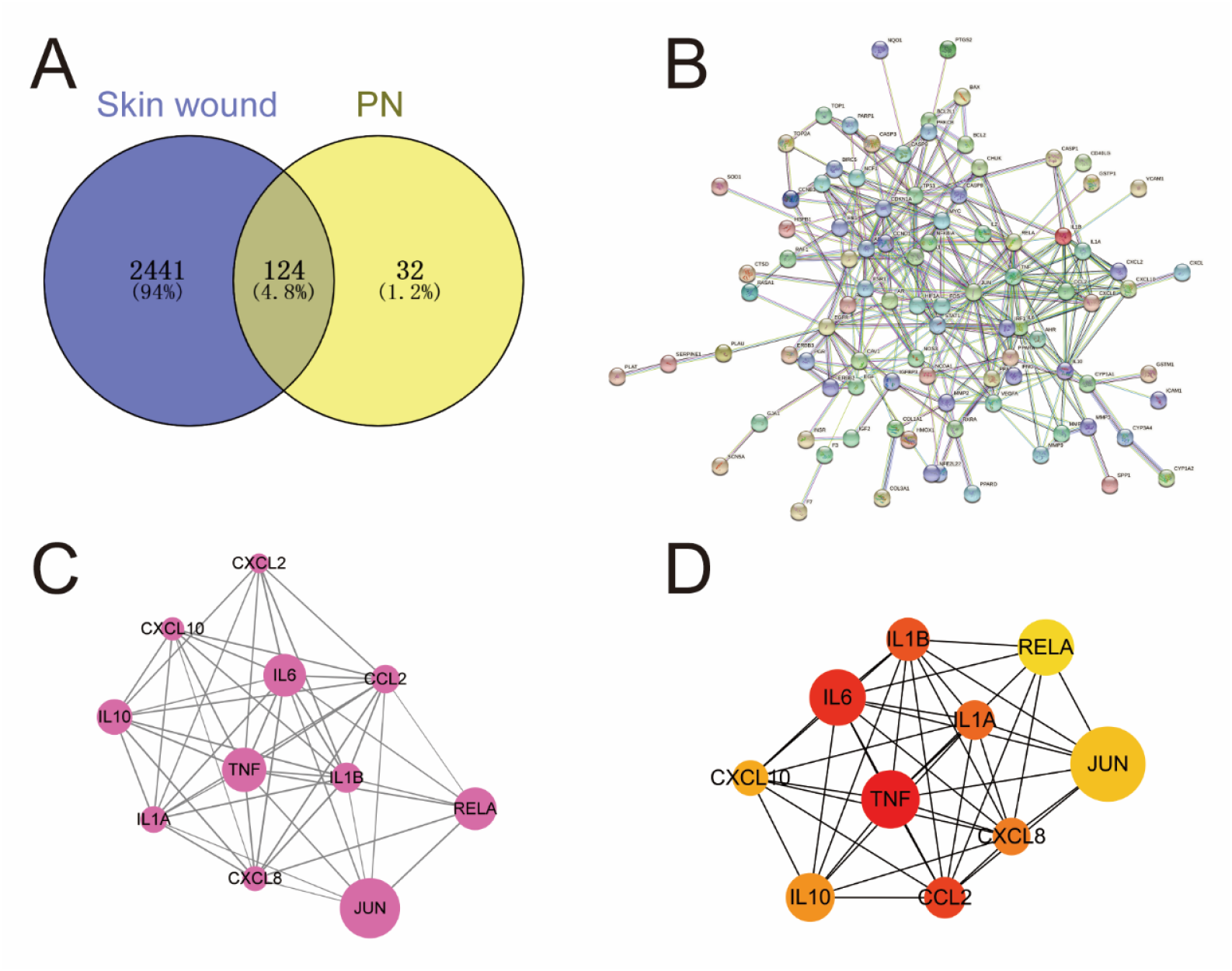
Common target screening and its interaction network; **PN:** Panax notoginseng**; A:** Intersection targets of panax notoginseng and skin trauma; **B:** PPI network of intersection targets of panax notoginseng and skin trauma; **C:** Major subnetworks (functional modules) in the target network. In the figure, node size is set according to node degree value, with larger nodes indicating greater importance. Edge thickness is set according to binding scores, with thicker edges indicating stronger PPI relationships; **D.** Top 10 key target genes ranked based on ECC in intersection targets, with redder colors indicating larger nodes and higher rankings.

### Intersection target function and pathway enrichment analysis

The intersection targets of PN and skin trauma were imported into the David database for GO and KEGG enrichment analysis. A threshold of P < 0.05 was set. A total of 712 GO annotation entries were obtained, including 562 biological processes (BP), 49 cell components (CC), and 101 molecular functions (MF). The top 10 GO entries were selected and plotted in ascending order of P value (**Fig 2A**). The GO enrichment results showed that biological processes such as positive regulation of RNA polymerase II promoter transcription, drug response, response to lipopolysaccharide, positive regulation of DNA template transcription, aging, response to estradiol, hypoxia response, positive regulation of nitric oxide biosynthesis process, positive regulation of gene expression, and negative regulation of apoptosis process all play important roles in the treatment of skin trauma with PN. Most of the core targets are located in the extracellular space, extracellular region, membrane rafts, caveolae, nucleus, cytoplasm, nuclear chromatin, and outer mitochondrial membrane.

**Fig 2.**
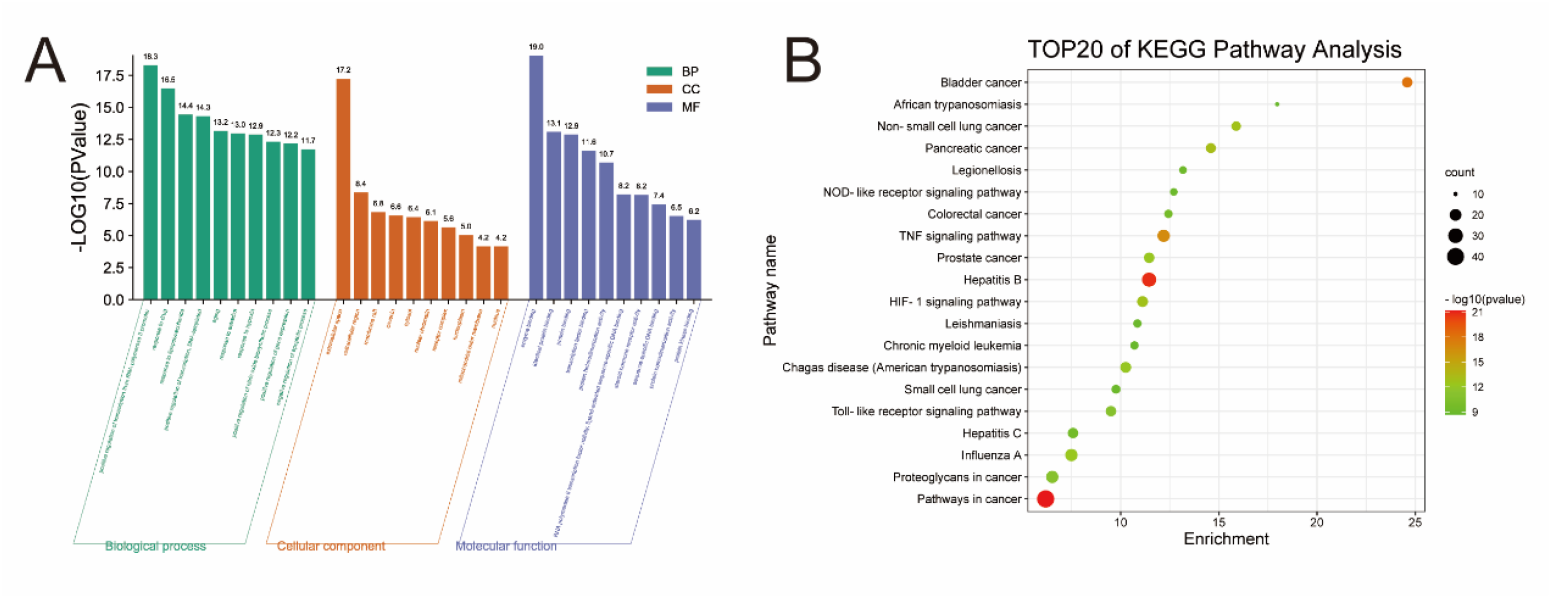
Functional enrichment analysis of intersection target genes. **A:** GO enrichment analysis (top 10)**; B:** KEGG pathway enrichment (top 20)

In the KEGG enrichment analysis, 115 signaling pathways were identified, and the top 20 entries were selected to create a bubble plot (**Fig 2B**). KEGG analysis revealed that the treatment of skin wounds with Panax notoginseng involves multiple signaling pathways, such as the cancer pathway, tumor necrosis factor signaling pathway, HIF-1 signaling pathway, Toll-like receptor signaling pathway, and NOD-like receptor signaling pathway. Furthermore, KEGG analysis also indicated that Panax notoginseng has potential therapeutic benefits for other diseases, including hepatitis B, American trypanosomiasis, leishmaniasis, hepatitis C, and chronic myeloid leukemia.

### Drug-component-target-pathway network

Using Cytoscape 3.8.2 software, a drug-component-target-pathway network was constructed based on the effective components of PN, intersecting target genes, and key signaling pathways (**Fig 3**).

**Fig 3.**
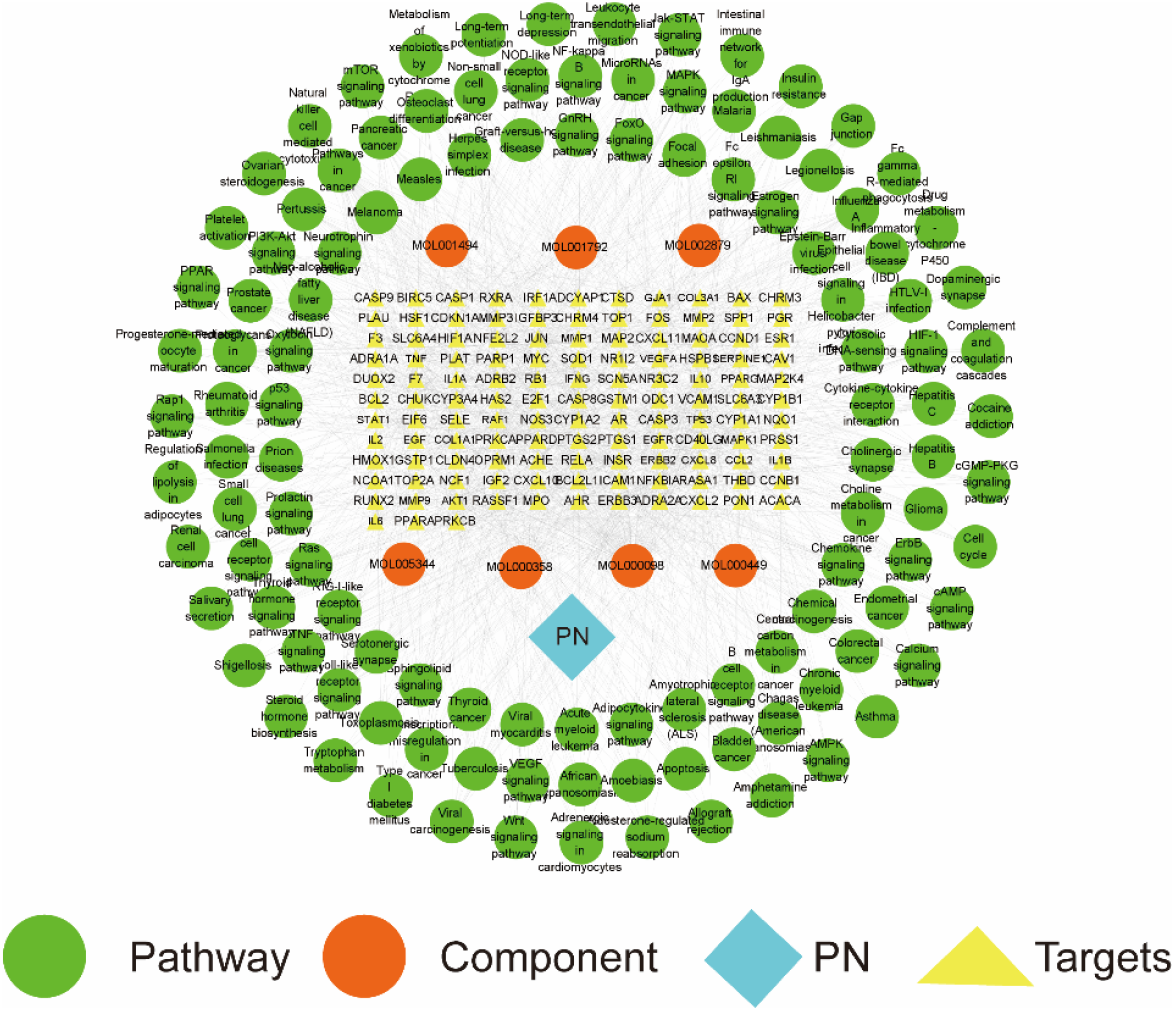
Drug-component-target-pathway network of PN in treating skin wound

### Measurement results of rat skin wound area

At 1d, 4d, and 7d after injury, the area of traumatized skin in both the PN group and the control group gradually decreased. At 4d and 7d, the area of skin wound in the PN group was smaller than that in the control group (P<0.05) (**Fig 4** and **Table 3**).

**Table 3.** Measurements of skin wound area in two groups at different time points.

| Group (n) | Wound Area (mm<sup>2</sup>) |  |  |
| --- | --- | --- | --- |
|  | 1d | 4d | 7d |
| NC (24) | 57.46±0.948 | 35.98±1.079 | 27.22±1.041 |
| PN (24) | 56.87±1.804 | 29.69±1.279<sup>Δ</sup> | 13.83±1.344<sup>Δ</sup> |

**Note:** Compared with the regular control group, <sup>Δ</sup>P< 0.05.

**Fig 4.**
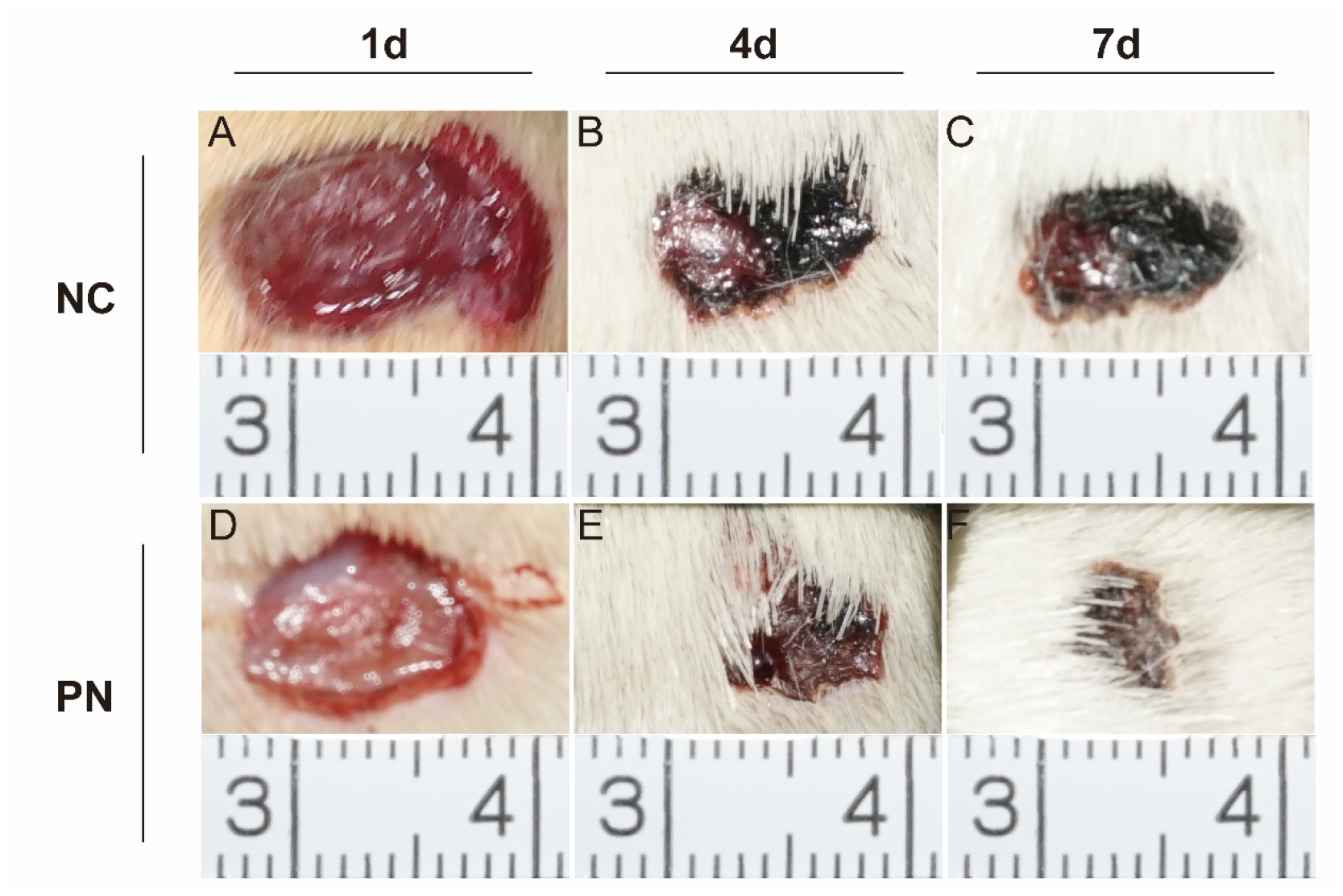
Skin trauma area. **NC:** regular control group**(A, B, C);PN:** Panax notoginseng group**(D, E, F); A:** Skin wound surface of NC 1 day after injury;**B:** Skin wound surface of NC 4 day after injury; **C:** Skin wound surface of NC 7 day after injury; **D:** Skin wound surface of PN 1 day after injury; **E:** Skin wound surface of PN 4 day after injury; **F:** Skin wound surface of PN 7 day after injury.

### HE staining results

One day after injury, under light microscopy, dilated and congested capillaries in the dermis layer of the injury area were observed in both groups, with a large number of red blood cells (indicated by red arrows) scattered in the tissue spaces and a small number of neutrophils (indicated by green arrows) infiltrating (**Fig 5-A1, A2, D1, D2**). Four days after injury, a large number of neutrophils (indicated by green arrows) and macrophages (indicated by white arrows) were observed infiltrating around the injury area in both groups, along with a significant number of fibroblasts (indicated by yellow arrows) and new capillaries (indicated by blue arrows) forming (**Fig 5-B1, B2, E1, E2**). Seven days after injury, in the PN group, the neutrophils (indicated by green arrows) and inflammatory cells around the injury area were significantly reduced compared to the control group, while a large number of fibroblasts (indicated by yellow arrows) proliferated and secreted collagen (**Fig 5-C1, C2, F1, F2**).

**Fig 5.**
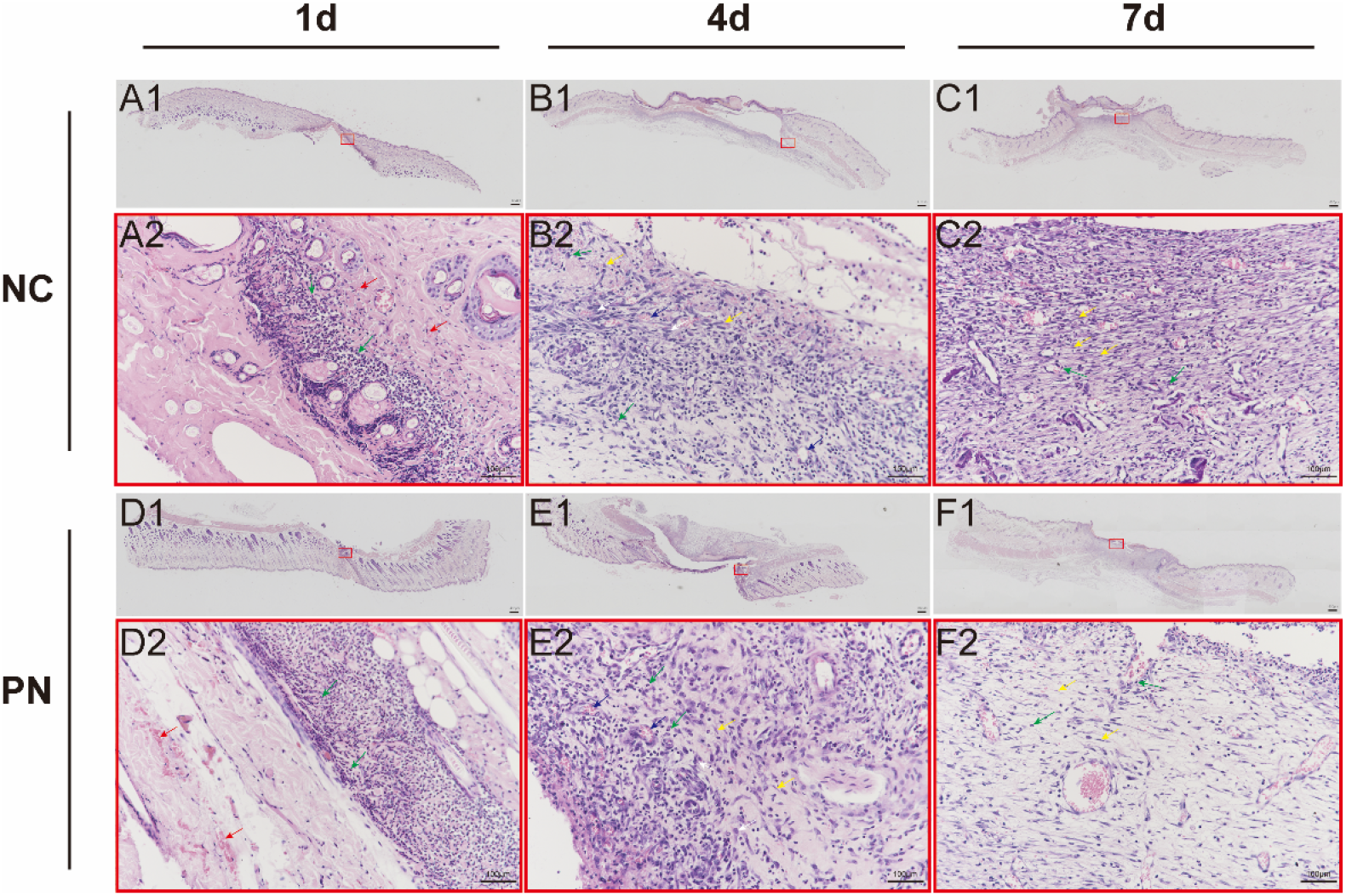
HE staining of skin wound areas; **NC:** regular control group;**PN:** Panax notoginseng group; **A1:** Skin tissue of NC 1 day after injury; **A2:** a partial enlarged view of A1**; B1:** Skin tissue of NC 4 day after injury; **B2:** a partial enlarged view of B1**; C1:** Skin tissue of NC 7 day after injury; **C2:** a partial enlarged view of C1**; D1:** Skin tissue of PN 1 day after injury; **D2:** a partial enlarged view of D1**; E1:** Skin tissue of PN 4 day after injury; **E2:** a partial enlarged view of E1**; F1:** Skin tissue of PN 7 day after injury; **F2:** a partial enlarged view of F1. Red blood cells are indicated with **red arrows;** Neutrophils are indicated with **green arrows;** Macrophages are indicated with **white arrows;** Fibroblasts are indicated with **yellow arrows;** The newly formed capillaries are indicated by **blue arrows**.

### IL-6 test results

Under light microscopy, IL-6-positive reactants were distributed in rat skin epidermal cells, hair follicles, neutrophils, macrophages, and fibroblasts, localized in the cytoplasm and matrix **(Fig 6 A(1,2)-F(1,2)**). The results of qRT-PCR and ELISA detection indicated that in the control group, the expression of IL-6 gradually increased from 1 to 7 days after injury; in the PN group, the expression of IL-6 reached its peak at 4 days, followed by a gradual decrease. Statistical analysis showed that at different time points, the expression intensity of IL-6 in the PN group was lower than that in the control group (P<0.01, **Fig 6G-H**).

**Fig 6.**
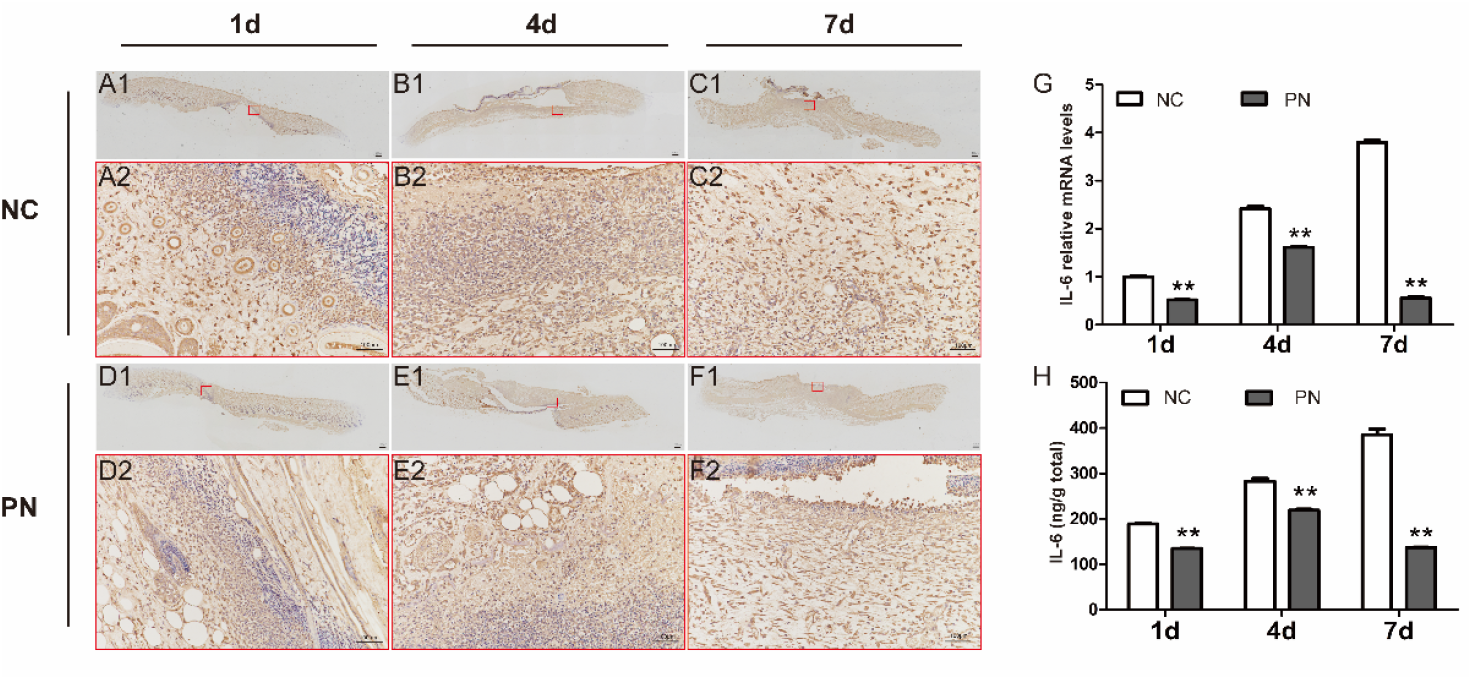
Detection results of IL-6 in skin wound area; **NC:** regular control group;**PN:** Panax notoginseng group; **A(1, 2)∼F(1, 2):** Immunohistochemical staining of IL-6 in skin wound area; **G:** relative mRNA expression level of IL-6; **H:** IL-6 protein expression level detected by ELISA; Comparison between PN and NC, * P<0.05; **P<0.01

### TNF-α detection results

Under light microscopy, TNF-α positive reactants were mainly distributed in the cytoplasm and matrix of rat skin epidermal cells, hair follicles, neutrophils, macrophages, and fibroblasts (**Fig 7A(1,2)-F(1,2)**). The qRT-PCR and ELISA results showed that after injury, the expression of TNF-α gradually increased in both the PN group and the control group. The statistical results showed that the expression intensity of TNF-α in the PN group was significantly lower than that in the control group at 4 and 7 days after injury (P<0.01, **Fig 7G-H**).

**Fig 7.**
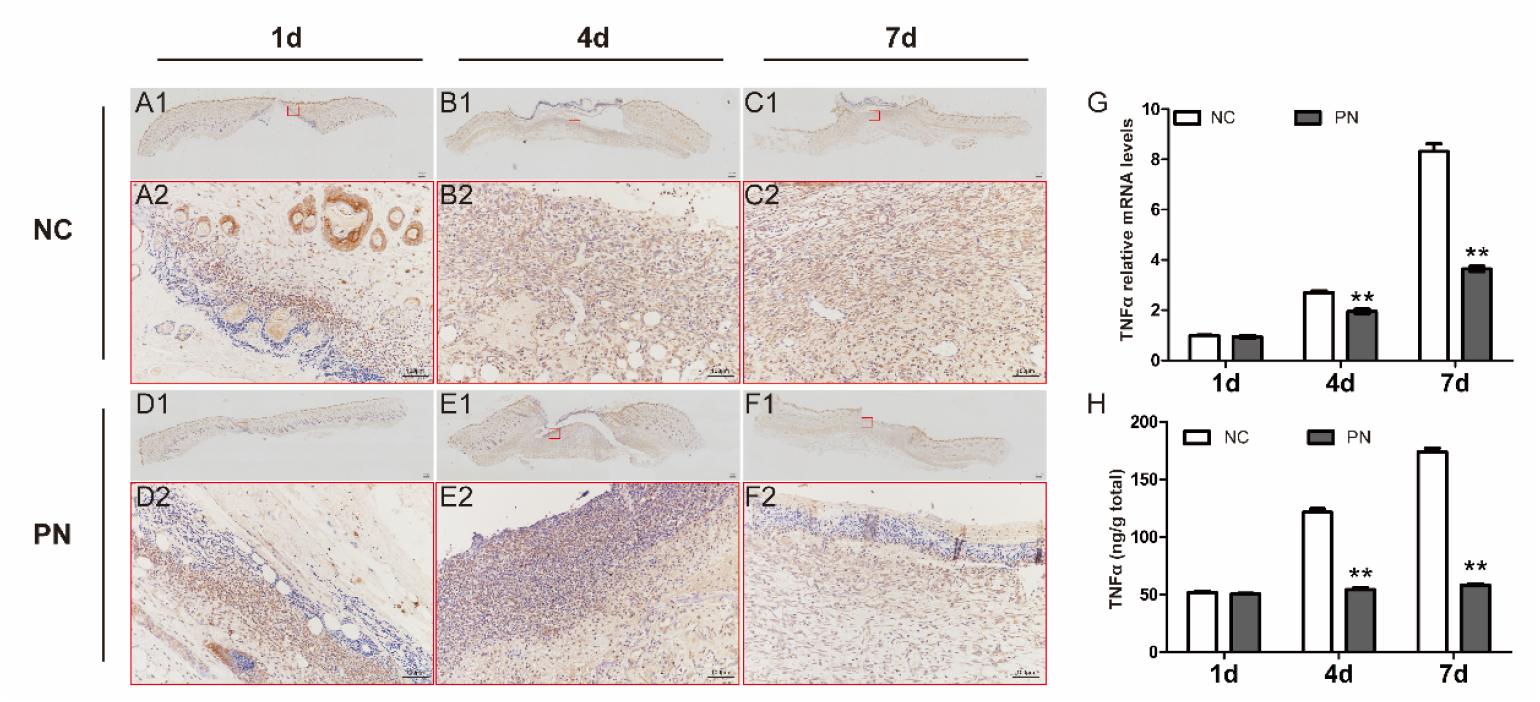
Detection results of TNF-α in skin wound area; **NC:** regular control group;**PN:** Panax notoginseng group; **A(1, 2)∼F(1, 2):** Immunohistochemical staining of TNF-α in skin wound area; **G:** relative mRNA expression level of TNF-α; **H:** TNF-α protein expression level detected by ELISA; Comparison between PN and NC, * P<0.05; **P<0.01

### IL-10 detection results

Under light microscopy, IL-10 positive reactants were mainly distributed in the cytoplasm and matrix of rat skin epidermal cells, infiltrating neutrophils, and macrophages. IL-10 expression was also observed in small amounts in some fibroblasts (**Fig 8A(1,2)-F(1,2)**). The results of qRT-PCR and ELISA detection showed that after injury, the expression of IL-10 gradually increased in both the PN group and the control group. The statistical results showed that the expression intensity of IL-10 in the PN group was significantly lower than that in the control group at 1 and 7 days after injury (P<0.01, **Fig 8G-H**).

**Fig 8.**
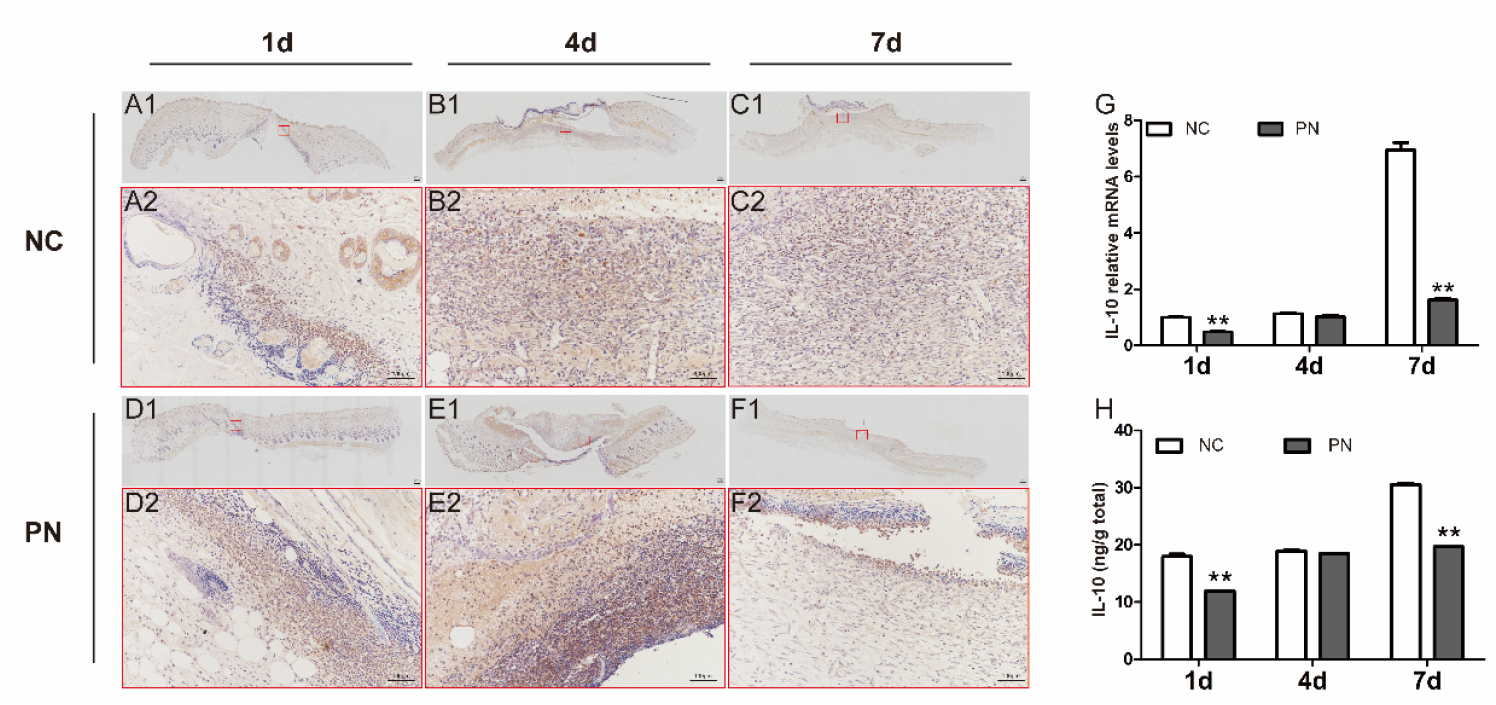
Detection results of IL-10 in skin wound area; **NC:** regular control group;**PN:** Panax notoginseng group; **A(1, 2)∼F(1, 2):** Immunohistochemical staining of IL-10 in skin wound area; **G:** relative mRNA expression level of IL-10; **H:** IL-10 protein expression level detected by ELISA; Comparison between PN and NC, * P<0.05; **P<0.01

## Discussion

### Exploring the Results of Network Pharmacology Analysis

This analysis shows that PN may affect the core metabolic pathways such as NF-κB signaling pathway, MAPK signaling pathway, JAK-STAT signaling pathway and chemokine signaling pathway through the regulatory network composed of TNF, IL-6, IL-10, CC-L2, IL-1B, IL-1A, RELA, CXCL-8, CXCL-10, JUN, etc., with multi-target and multi-channel coordination and multiple regulation. Among them, TNF, IL-6 and IL-10 are the core target combinations of PN, which constitute the “pro-inflammatory-anti-inflammatory” control axis of inflammatory reaction. They can form a complex positive and negative feedback loop through autocrine and paracrine, which runs through three stages of skin wound healing: hemostasis, inflammation, proliferation and remodeling, and cooperatively regulate the skin wound healing process [11, 12].

Related studies have shown that TNF is the core factor to initiate early inflammation and can directly induce IL-6 gene transcription by activating NF-κB and MAPK pathways [13, 14]; After IL-6 is secreted, it can further amplify the inflammatory response through the JAK-STAT3 pathway, form a TNF-IL6 positive feedback loop, promote neutrophil infiltration and increase vascular permeability [15, 16]. This loop can clear pathogens and necrotic tissue in the early stage of trauma, but excessive activation can lead to uncontrolled inflammation, triggering tissue edema and fibrosis [17, 18]. IL-10 is a classic anti-inflammatory cytokine, which can inhibit the overexpression of TNF/IL-6 in two ways: one is to directly inhibit the NF-κB pathway, reduce RELA into the nucleus, and block the transcription of TNF and IL-6 [19-21]; The second is to activate STAT3, induce the expression of anti-inflammatory genes (such as SOCS3), negatively regulate the JAK-STAT pathway, and inhibit IL6 signaling [22, 23]. In addition, IL-10 can also inhibit the polarization of macrophages to a pro-inflammatory phenotype (M1 type) and promote their conversion to an anti-inflammatory repair phenotype (M2 type), thereby reducing TNF and IL-6 secretion and increasing growth factors (such as TGF-β) release [24, 25]. Under physiological conditions, the inflammatory response shows temporal changes: in the early stage (0-3 days after trauma), TNF and IL-6 are mainly high expressed, and immune cell recruitment and tissue debridement are initiated; IL10 expression gradually increased in the middle stage (3-7 days), which inhibited excessive inflammation and promoted the transition from inflammation to repair stage [26, 27]; If IL-10 expression is insufficient or TNF/IL-6 is over-activated, inflammation will be prolonged [28, 29].

It is suggested that PN may regulate the balance of pro-inflammatory and anti-inflammatory and the time sequence transition of inflammation-repair by targeting the core factors such as TNF, IL-6 and IL-10 and their rich NF-κB and MAPK pathways. Among them, TNF/IL6 constitutes a positive feedback loop to promote inflammation and start the early immune response after trauma. As a negative regulatory factor, IL-10 inhibits inflammation from getting out of control and promotes repair. The cross-talk among them is the core axis of skin wound healing. PN may realize the synergistic effect of “anti-inflammatory-repair” by regulating the network in two directions.

### Exploration of experimental results

This validation experiment revealed the temporal and spatial expression rules of IL-6, TNF-α and IL-10 in skin wound healing from three aspects: protein localization, gene transcription and protein secretion, and also clarified the mechanism of PN in optimizing the balance between inflammation and repair by reshaping the expression sequence and intensity of these three core cytokines. The experimental results are highly consistent with the aforementioned “pro-inflammatory-anti-inflammatory control axis” and pathological features, which provides direct molecular biological evidence for PN to promote wound healing.

The results of this experiment showed that there was no significant difference in trauma area between the two groups on the first day after trauma (P > 0.05). A small number of neutrophils, inflammatory cell infiltration, dermal telangiectasia and hyperemia appeared in the skin wound area of both groups. This may stem from the physiological initial response of wound healing, that is, immediately after trauma, damaged cells release “initial pro-inflammatory factors” such as TNF, IL1B, and IL1A, activate the RELA/JUN pathway, induce the expression of chemokines such as CXCL8, recruit a small number of neutrophils to the wound, and at the same time start vasodilation and congestion to deliver immune cells and nutrients [30-34]. At this stage, Panax notoginseng showed no significant effect, suggesting that Panax notoginseng did not inhibit the initiation of inflammation. At 4d and 7d after trauma, the wound area of Panax notoginseng group was significantly smaller than that of the control group (P < 0.05). The results of HE staining showed that a large number of neutrophils and macrophages infiltrated around the injured area in the control group (4 days after trauma, that is, the early stage of proliferation and repair), which suggested that the skin wound in the control group was still dominated by inflammatory reaction at this stage, and fibroblast proliferation and collagen synthesis had not been completely started. On the 7th day after trauma, the inflammatory cells around the injured area in Panax notoginseng group (middle stage of proliferation and repair period) decreased significantly, and fibroblasts proliferated in large quantities and secreted collagen. This indicates that the Panax notoginseng group may accelerate the transition of “inflammation-repair” by inhibiting excessive inflammation in advance, so that fibroblasts can enter the proliferation cycle earlier, and then accelerate the wound contraction rate [35].

The results of immunohistochemical staining showed that IL-6 and TNF-α were located in the cytoplasm and matrix of epidermal cells, hair follicles, neutrophils, macrophages and fibroblasts, suggesting that the synthesis and function of these two pro-inflammatory factors run through the core cell group of wound healing. Related studies have shown that IL-6 and TNF-α secreted by epidermal cells/hair follicles can start the proliferation and migration of local epithelial cells and participate in the re-epithelization process of wounds [36-38]; IL-6 and TNF-α secreted by neutrophils/macrophages as the core immune cells of inflammatory reaction can amplify the inflammatory signal, recruit more immune cells to the wound surface, and regulate the phenotypic polarization of macrophages [3, 39, 40]; IL-6 and TNF-α synthesized by fibroblasts can regulate their own proliferation and collagen synthesis by autocrine/paracrine, but their over-expression will inhibit the repair function of fibroblasts [41, 42]. The results of this experiment show that IL-10 is mainly localized in the cytoplasm and matrix of epidermal cells, neutrophils and macrophages, and only a small amount is expressed in fibroblasts, which suggests that the core target of IL-10 is mainly focused on immune cells. Related research shows that macrophages are the main source of IL-10, and the secreted IL-10 can reverse the polarization of macrophages from M1 type (pro-inflammatory) to M2 type (anti-inflammatory repair) [43]. IL-10 secreted by epidermal cells can maintain the local immune homeostasis of wound surface and reduce the damage of inflammation to epithelial regeneration [44, 45]; The low expression of IL-10 in fibroblasts can moderately inhibit excessive collagen deposition and avoid scar hyperplasia [46, 47].

This study shows that in the early stage (1-3 days) after skin trauma, PN can maintain moderate expression of TNF-α and low expression of IL-10, ensuring the initiation and debridement of inflammation [48]; At the same time, IL-6 was slightly inhibited to avoid premature amplification of inflammation [49]. In the middle stage (4d) after skin trauma, PN makes IL-6 fall back in time after reaching its peak, which significantly inhibits the over-activation of TNF-α and starts the inflammation-repair transition [50, 51]; At this time, moderate expression of IL-10 promoted the polarization of macrophages to M2 type [52]. In the late stage of skin trauma (7 days): PN continued to inhibit TNF-α/IL-6, the inflammatory reaction basically subsided, and the low expression of IL-10 could maintain a steady state; Fibroblasts proliferate in low inflammatory environment, secrete collagen and accelerate wound healing [53, 54]. However, the control group was caught in a vicious circle of \over-inflammation-passive anti-inflammation\ due to the continuous high expression of TNF-α/IL-6 and the compensatory increase of IL-10, which eventually led to delayed healing [55]. It is suggested that PN has a sequential advantage in the treatment of skin wounds. By reshaping the expression sequence and intensity balance of TNF-α/IL-6/IL-10, an ideal healing microenvironment of “efficient debridement, timely anti-inflammatory and orderly repair” can be constructed, which can shorten the critical time window of wound healing and promote the rapid healing of skin wounds.

## Conclusions

To sum up, PN shows a significant effect on promoting healing in the treatment of skin wounds. It has the characteristics of multi-components, multi-targets, multi-pathway synergy, and multiple regulatory pathways. Its core mechanism lies in precisely regulating the local inflammation-repair balance of wounds. By targeting the regulatory network composed of core cytokines such as TNF-α, IL-6, and IL-10, PN can achieve timing and precise intervention: early ensuring the initiation of physiological inflammation to complete wound debridement, effectively inhibiting TNF-α and IL-6 overexpression in the middle and late stages, avoiding prolonged inflammation and secondary tissue damage, and at the same time promoting the transformation of IL-6 from pro-inflammatory to pro-repair function, and promoting the polarization of macrophages to anti-inflammatory repair phenotype. This series of regulation is finally reflected in the reduction of inflammatory cell infiltration, massive proliferation of fibroblasts and orderly secretion of collagen at the pathological level, while phenotypic significantly accelerates wound contraction and shortens healing time. In short, PN provides a therapeutic effect of “efficient debridement-orderly repair-low scar risk” for skin wounds by remodeling the dynamic balance of cytokine network and optimizing the time sequence process of “inflammation initiation-repair conversion-tissue remodeling”. It is one of the ideal natural medicines to regulate skin wound healing.

## Authors’ contributions

**Data curation:** Jun Liu, Jun-cen Li.

**Formal analysis:** Hua-man Geng, Gen-ke Li.

**Funding acquisition:** Zhi-hong Zhang.

**Methodology:** Chen Jin, Jie Luo.

**Resources:** Zhi-hong Zhang.

**Writing – review & editing:** Yi-bin Li, Qing-lin Li.

